# A contribution to the larval amphibian microbiome: characterization of bacterial microbiome of *Ichthyophis bannanicus* (Order: Gymnophiona) and comparison with the other two amphibian orders

**DOI:** 10.1101/2021.09.20.461075

**Authors:** Amrapali Prithvisingh Rajput, Shipeng Zhou, Madhava Meegaskumbura

**Affiliations:** Eco.Evo.Devo Lab-Group, Guangxi Key Laboratory of Forest Ecology and Conservation, College of Forestry, Guangxi University, Nanning, Guangxi, People’s Republic of China

**Keywords:** amphibia, gut-skin axis, *Ichthyophis bannanicus*, metabarcoding, 16S rRNA

## Abstract

It is known that animal-associated microbiomes form indispensable relationships with hosts and are responsible for many functions important for host-survival. Next-gen driven approaches documenting the remarkable diversity of microbiomes have burgeoned, with amphibians too, benefiting from such treatments. The microbiome of Gymnophiona (caecilians), one of the three amphibian orders, constituting of 3% of amphibians, however, remains almost unknown. The present study aims to address this knowledge gap through analysis of the microbiome of *Ichthyophis bannanicus*. As these caecilian larvae are aquatic and hence exposed to a greater propensity for bacterial microbiomic interchange, we hypothesize that bacterial phyla would overlap between gut and skin. Further, from the host-specificity patterns observed in other vertebrate taxa, we hypothesize that Gymnophiona have different dominant gut bacterial microbiomes at a higher taxonomic level when compared to the larvae of the other two amphibian orders (Anura and Caudata). We used 16S rRNA gene amplicon sequencing based on Illumina Nova sequencing platform to characterize and compare the gut (represented by faecal samples) and skin microbiome of *I. bannanicus* larvae (*N* = 13), a species distributed across South-East-Asia and the only caecilian species occurring in China. We compared our gut microbiome results with published anuran and caudate larval microbiomes. For *I. bannanicus*, a total of 4,053 operational taxonomic units (OTU) across 13 samples were detected. Alpha-diversity indices were significant between gut and skin samples. Non-metric multidimensional scaling analysis suggest that gut and skin samples each contained a distinct microbiome at OTU level. We record significant differences between the bacterial phyla of gut and skin samples in larvae of *I. bannanicus*. The study provides an overview of gut and skin bacterial microbiomes of a caecilian, while highlighting the major differences between larval microbiomes of the three amphibian orders. We find a partial overlap of gut bacterial microbiomes at phylum level for the three orders; however, the relative abundance of the dominant phyla is distinct. The skin and gut microbiomes are distinct with little overlap of species, highlighting that gut-skin axis is weak. This in turn suggests that many of the microbial species on skin and gut are functionally specialized to those locations. We also show that the skin microbiome is more diverse than the gut microbiome at species level; however, a reason for this could be a portion of the gut microbiome not being represented in faecal samples. These first microbiome information from a caecilian lay the foundation for comparative studies of the three amphibian orders.

## INTRODUCTION

The bacterial microbiome (BM), shaped by the life-history strategies and phylogenetic relationships of hosts, is now known to be closely associated with host well-being (*Hanning & Diaz-Sanchez, 2015; Davenport et al., 2017*). Among vertebrates, amphibians are highlighted as having complex life cycles, reproductive mode diversity, occupation of a diversity of habitats, a multitude of food preferences (*Wilbur, 1980; Duellman & Trueb, 1986; McDiarmid & Altig, 1999*), and hence are expected to show microbiome assemblages reflective of these life history strategies. Among recent amphibian BMs studied (between 2005-2020), a lion’s share pertains to anurans (frogs and toads, 89%), with urodeles (salamanders and newts, 11%) constituting the remainder. An entire order, Gymnophiona (caecilians), remains unstudied (*Wiggins et al., 2011; Mashoof et al., 2013; Kohl et al., 2013, 2014, 2015; Bletz et al., 2016; Chang et al., 2016; Vences et al., 2016; Weng et al., 2016; Zhang et al., 2016; Knutie et al., 2017, Warne et al., 2017; Demircan et al., 2018; Huang et al., 2018; Lyra et al., 2018; Mu et al., 2018; Zhang et al., 2018; Tong et al., 2019a, b; Warne et al., 2019; Ya et al., 2019; Long et al., 2020; Xu et al., 2020; Zhang et al., 2020*). This pattern of study emphasis of host BMs appears to reflect the diversity of species of the three orders – of a total of 8,176 amphibian species, 7,212 are Anura (frogs and toads, 88%) and 750 are Caudata (newts and salamanders, 9%), while only 214 are Gymnophiona (caecilians, 3%) (*Amphibiaweb, 2021*).

Gymnophiona occur across much of the wet tropics and some subtropical regions apart from Madagascar, Australia, South-East Asia and East of Wallace’s line (*Taylor, 1968*). *Ichthyophis bannanicus* is the only caecilian species found in China (*Wang et al., 2015*). These legless amphibians that lead a fossorial or aquatic life, are secretive and difficult to locate. Known as Banna caecilian, the species ranges across Southern China. It has an aquatic larval stage of about two years (*Li et al., 2010*) and a fossorial post-metamorphic stage during which it rarely enters water bodies (*Meng & Li, 2006*). Almost nothing is known of the BMs of larval or the adult caecilians.

Gut microbial diversity is known to be influenced by the growth environment, developmental stages, and health conditions of the host (*Tong et al., 2019a: Long et al., 2020*). As amphibians metamorphose, their symbiotic gut BM also alters, providing nutritional needs depending on food habits and habitat of the relevant life-history stage (*Zhang et al., 2020*). The role of digestion and nutrient uptake by the BM ranges from the breakdown of various complex food sources into easily absorbable nutrients, to the manufacturing of secondary metabolites. Studies on *Lithobates pipiens*, the Northern leopard frog, suggest that the gut microbial diversity of tadpoles is similar to that of fishes, while the adult gut microbiota is similar to that of terrestrial vertebrates (*Kohl et al., 2013*). Hence, the biphasic life histories of aquatic larval stages and semi-terrestrial adults of amphibians (*Wilbur, 1980*) makes amphibians an ideal system to study the gastrointestinal bacteria and their role in enhancing host’s life processes.

Furthermore, amphibian skin, by providing a moist respiratory surface, is suitable for rich communities of microorganisms, both beneficial and detrimental to the host. However, the skin of an amphibian also has poison glands that discourage certain microbes. It is known, however, that the amphibian skin microbiome plays a vital role as symbionts, protecting their hosts against disease (*Rebollar et al., 2020*). Symbiotic skin bacteria may provide resistance to pathogens either by producing metabolites that directly impede pathogen growth, or by stimulating the host immune system (*Bletz et al., 2013*). As caecilian larvae are aquatic, the skin and gut may host similar microflora. Thus, the gut-skin axis is assumed to have beneficial roles by protecting the host species from diseases while helping in the re-colonization of essential BMs.

Here we focus on Gymnophiona, represented by the larvae of *I. bannanicus*. Our objective is to study the BM of gut (fecal samples were used as a proxy for the gut) and skin. As the larvae are aquatic, the skin is inhabited by microbes present in waterbodies. Caecilians feed on organisms inhabiting waterbodies; at the same time, water enters the digestive system while feeding, but the stomach acts as a filter for the progression of certain bacteria to the lower gut. Thus, we hypothesize that some of the bacterial phyla would overlap between gut and skin. As caecilian larvae have a carnivorous lifestyle, as opposed to the larvae of anurans and caudates, we hypothesize that the gut microbiota of *I. bannanicus* would be distinct at a higher taxonomic level.

Since the skin of larval caecilians sheds periodically, and the larvae live in an aquatic medium, we hypothesize that skin is recolonized from bacteria living in the gut through fecal transmission. Hence, we expect the skin BM to be less diverse than the gut BM. We investigate the BM using high-throughput sequencing of the bacterial 16S rRNA gene fragment.

Our study reveals that although the bacterial diversity of the gut partially overlaps the larvae of the three amphibian orders, the relative abundance of dominant phyla remains distinct. However, we observed that the skin BM was more diverse than that of the gut, we discuss an explanation for this.

## MATERIALS AND METHODS

*This study was carried out in accordance with the approval of Institutional Animal Care and Use Committee of Guangxi University (GXU), Nanning-China. Animal procedures were by GXU approval document (#GXU2019-071)*.

### Sample Collection

Larvae of *I. bannanicus* were obtained from a pet-market and maintained in the laboratory. They were reared in plastic boxes consisting of dechlorinated water (3 cm, height). The water was renewed every two days. Caecilians were reared on frozen blood worms. Sample collection for gut samples (*N* = 13) was carried out by placing the test subjects temporarily in sterile water. As soon as the caecilian defecated, the sample was collected using a sterile dropper. Skin swabs were collected immediately before placing the larvae in sterile water. Vials containing samples were immediately frozen at −86 °C until DNA extraction. All samples from the test subjects were collected at larval development stage 37 (*Dunker et al., 2000*). Morphometric measurements (i.e., body weight, gm; body width, mm; and total length, mm) were recorded for each individual (Table S1). Sterile conditions were maintained throughout the procedures to prevent contamination of samples.

### DNA Extraction

Bacterial genomic DNA was extracted from larval gut and skin according to the manufacturer’s protocol, with the aid of Power Soil kits and DNA Tissue-Blood kits (QIAGEN, Hilden Germany). The robustness of the DNA was visually monitored using 1.0% agarose gel electrophoresis and quantified using a Qubit and NanoDrop. Hyper-variable regions of the 16S rRNA gene (V3-V4) were PCR-amplified from genomic DNA using the bacteria-specific universal barcode-primers 515F and 806R. All polymerase chain reactions were performed using 15 μL of Phusion ® High-Fidelity PCR Master Mix, 0.2 μM of each forward and reverse primer and 10 ng of DNA template. Thermal cycling conditions were as follows: initial denaturation at 98 °C for 1 min, followed by 30 cycles of denaturation at 98 °C for 10 s, annealing at 50 °C for 30 s, elongation at 72 °C for 30 s and final extension at 72 °C for 5 min.

### Illumina Library Preparation

Amplicons of each PCR sample were extracted by mixing same volume of 1X loading buffer containing SYB green with PCR products and further electrophoresis was operated on 2% agarose gel for detection. PCR products were mixed in equidensity ratios. The PCR products were further purified using Qiagen Gel Extraction Kits. The sequencing libraries were generated using TruSeq® DNA PCR-Free Sample Preparation Kit following the manufacturer’s protocol, followed by the addition of index codes. Evaluation of library quality was performed on a Qubit@ 2.0 Flurometer and Agilent Bioanalyzer 2100 system. Further, the library was sequenced on an Illumina NovaSeq platform and 250 bp paired-end reads were generated. Paired-end reads were assigned to the samples based on their unique barcode, truncated by cutting off the barcode and primer sequence. The paired-end reads were further merged using FLASH and the sequences spliced. The quality filtering was performed according to the QIIME (Version 1.9.1) quality control protocol (*Caporaso et al., 2010*). The tags were compared with reference database using UCHIME algorithm to detect chimera sequences, and effective tags were thus obtained.

### 16S rRNA Gene Sequence Analysis

Sequence analysis was performed using Uparse software. Sequences with approximately 97% similarity were assigned to the same OTUs (*Rideout et al., 2014*). A representative sequence for each OTU was screened for further annotation. For each representative sequence, the SILVA database, which uses the Mothur algorithm, was used to annotate the taxonomic information. To study phylogenetic relationships of different OTUs and the differences amongst the dominant species in different samples, multiple sequence alignment was conducted using MUSCLE software. OTUs abundance information was further normalized using the standard sequence number corresponding to the sample with least sequences. The SILVA 123 database was implied for taxonomic assignment. Reference sequences in the SILVA 123 database were initially trimmed to hypervariable region (V3-V4) with 515F-806R universal primers used in the PCR. Taxonomic assignments were carried out using UCLUST with a minimum confidence threshold of 80% (*Edgar, 2010*). Libraries containing at least 1,000 reads were used for analysis, and sub-OTU relative abundance values were calculated by transformation to library read depth. We used circled legend and bar plot to show the microbial composition of each sample using R (4.1.1, “amplicon” package; Liu *et al*., 2015).

### Data Availability

All the raw metagenomic data is available through the National Center for Biotechnology (NCBI) with the BioProject accession number PRJNA764182.

### Data Analysis

Alpha and beta diversity were calculated using QIIME 1.9.1 version. Alpha diversity indices were compared using Wilcox rank-sum test, to estimate community diversity indices (Shannon; Simpson) and community richness indices (Abundance-based coverage estimator-ACE; Chao1). Beta diversity was calculated with unweighed UniFrac, weighed UniFrac and Bray-Curtis metrics (*Lozupone & Knight, 2005*). To estimate the gut and skin bacterial diversity we used “ggplot2” and “ggsignif” packages in R (4.1.1 version).

The Bray-Curtis distance for abundance was used to generate non-metric multidimensional scaling (NMDS) to visualize beta diversity patterns and reflect the inter and intra group differences using R (4.11, “vegan” and “ggplot2” packages. To test if the group dispersion (gut/skin), one way of similarity (ANOSIM) was performed based on OTUs. To investigate differences in the microbial community structure between the gut and skin samples, unifrac distance across each sample was implemented to generate UPGMA trees (unweighed pair-group method with arithmetic mean) using R (“hclust”). The difference between gut and skin microbial diversity at phylum level was analyzed using t-test.

Linear discriminant analysis effect size (LEfSe) was used to identify whether sub-OTUs significantly differ amongst gut and skin samples (*Segata et al., 2011*). The threshold in Kruskal-Wallis test amongst groups was considered to be statistically significant at *P* < 0.05. The taxa with a log LDA score (Linear discriminant analysis) more than four orders of magnitude were considered. Network analysis was conducted to show the co-occurrence and co-exclusion relationships among the abundances of clades in the gut samples *I. bannanicus* larvae.

## RESULTS

By splicing reads (sequencing based on the Illumina Nova sequencing platform by constructing a PCR-free library, followed by Paired-End sequencing), an average of 68,901 tags were measured per sample, and 66,100 effective data were obtained after quality control. The effective data volume of quality control reached 60,598, and the effective rate of quality control reached 87.98%. The read depths of the libraries were sufficient to capture the BM in both gut and skin, given that all libraries reached saturation in the rarefaction curve (Fig. 1A; Table 1). The rank abundance curve directly reflected the richness of bacterial communities in the samples (Fig. 1B). The BM in each of the gut and skin samples reached saturation in the rarefaction curve (Fig. S1). The rank abundance curve directly reflected the richness of bacterial communities in each of the gut and skin samples (Fig. S1).

**Figure 1.**
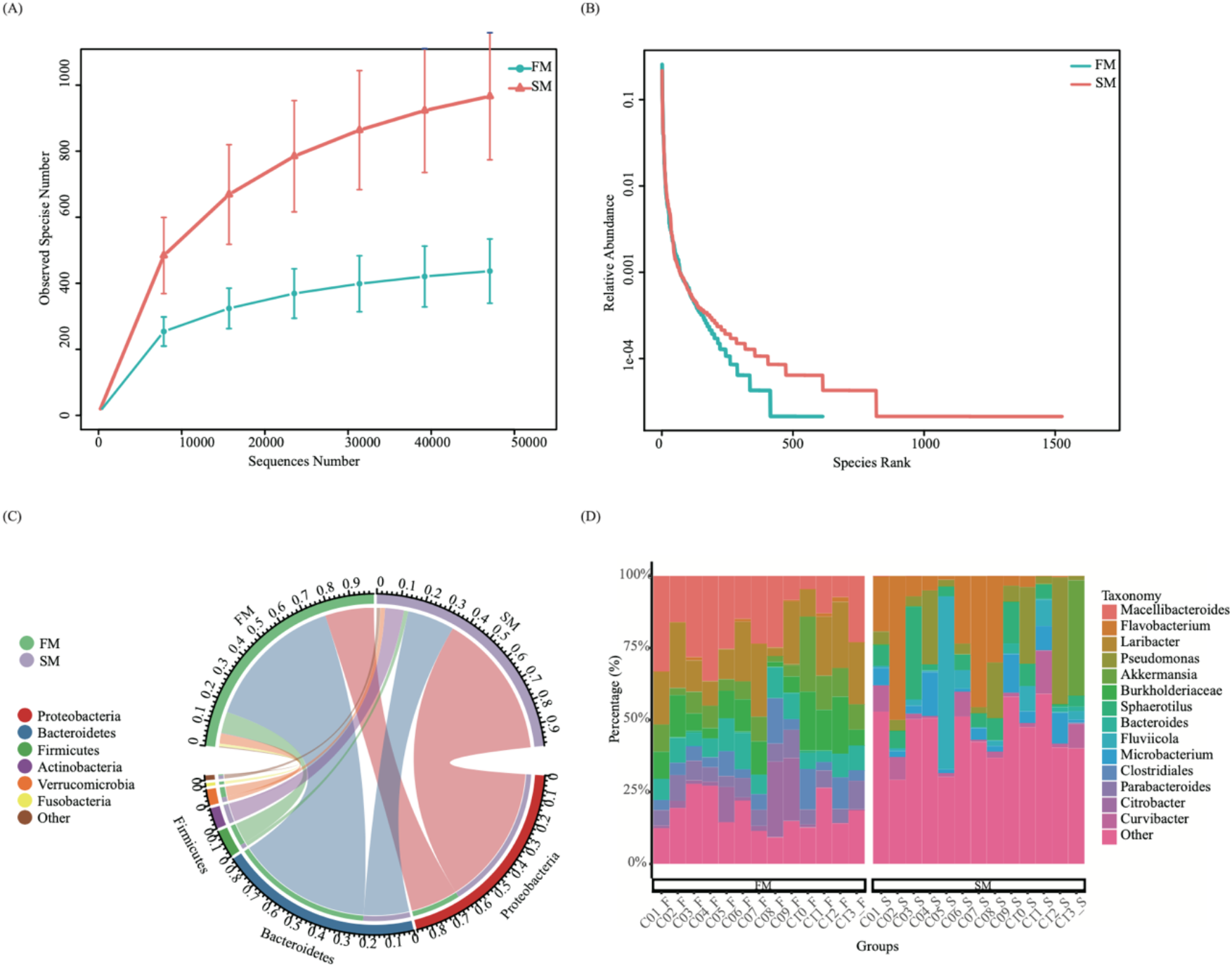
(A) The rarefaction analysis of the microbe species from gut (FM) and skin (SM) of *I. bannanicus* larvae; (B) The rank abundance curve analysis of the microbe species from gut (FM) and skin (SM) of *I. bannanicus* larvae; (C) Circular layout: Phylum level distribution of bacterial taxa of gut (FM) and skin (SM) in *I. bannanicus* larvae; (D) Distribution of bacterial taxa of gut (C01_F-C13_F) and skin (C01_S-C13_S) in *I. bannanicus* larvae genus level.

**Table 1.** The microbial diversity index including Chao1, ACE, Simpson and Shannon of 16S rRNA sequence library from 13 individuals of *Ichthyophis bannanicus* at a given rarefied depth.

| Diversity index | Gut sample | Skin sample |
| --- | --- | --- |
| Observed species | 436 | 966 |
| Shannon | 4.453 | 4.717 |
| Simpson | 0.874 | 0.836 |
| ACE | 491.046 | 1096.715 |
| Chao1 | 505.725 | 1141.309 |
| Goods coverage | 0.998 | 0.995 |
| PD whole tree | 52.002 | 121.213 |

The sequences were classified into 4,053 operational taxonomic units based on 97% sequence identity using QIIME 1.9.1 version (*Caporaso et al., 2010*). A total of 1,429 (35.26%) genera were identified when OTU sequences were tallied against the Silva123 database and annotated for taxon identity. The unique and commonly shared OTUs between the samples showed that, the highest frequency of OTUs were observed on skin samples when compared to the gut samples (Table S2).

### Bacterial abundance in gut and skin samples of *I. bannanicus*

The composition of the BM of gut samples in *I. bannanicus* was analyzed at six classification levels. At phylum level (Fig. 1C) Bacteriodetes, Proteobacteria, Firmicutes and Verrucomicrobia were dominant, accounting for 62.32%, 21.70%, 9.45% and 4.50% of OTUs respectively. Bacteroidia, Gammaproteobacteria and Clostridia were the dominant classes, which accounted for 62.32%, 20.79% and 8.38% of OTUs, respectively (Fig. S2A). At order level Bacteroidales (61.48%), unidentified_Gammaproteobacteria (17.17%) and Verrucomicrobiales (4.5%) were dominant (Fig. S2B). Members of Tannerellaceae (15.35%), unidentified_Gammaproteobacteria (9.00%) and Burkholderiaceae (7.17%) were dominant at family level (Fig. S2C). *Macellibacteroides* (11.44%) and *Laribacter* (8.98%) were the dominant genera (Fig. 1D). *Akkermansia glycaniphila* (3.96%), *Alcaligenaceae bacterium* BL-169 (3.54%) and *Bacteroides neonati* (0.97%) were dominant species in the gut samples (Fig. S2D).

The composition of BMs at six classification levels were analyzed for skin samples of *I. bannanicus*. At phylum level (Fig. 1C), Proteobacteria (64.49%), Bacteriodetes (20. 93%), Actinobacteria (8.36%) and Verrucomicrobia (2.09%) were dominant. At class level (Fig. S2A). Gammaproteobacteria and Bacteroidia were dominant, accounting for 57.93% and 20.88%, respectively. Unidentified_Gammaproteobacteria, Flavobacteriales and Pseudomonadales were the dominant orders (Fig. S2B) constituting of 41.85%, 14.63% and 9.36% of OTUs, respectively. At family level, members (Fig. S2C) of Burkholderiaceae (38.69%), Flavobacteriaceae (10.38%) and Puseudomonadaceae (8.15%) were dominant. *Flavobacterium* and *Pseudomonas* were dominant members at genus (Fig. 1D) level, consisting of 10.28% and 8.14% of OTUs, respectively. Dominant species (Fig. S2D) were *Microbacterium oxydans* (3.73%), *Sanguibacter inulinus* (1.51%) and *Klebsiella pneumoniae* (1.02%).

### Alpha and beta diversity

Comparison of alpha diversity indices were significant between gut and skin samples that were detected using the Wilcox test. The ACE (Fig. 2A) and Shannon index (Fig. 2B) of *I. bannanicus* skin microbiota samples were significantly higher than that of the gut microbiota samples. Both community richness and community diversity in *I. bannanicus* were distinct between gut and skin samples (Fig. S3).

**Figure 2.**
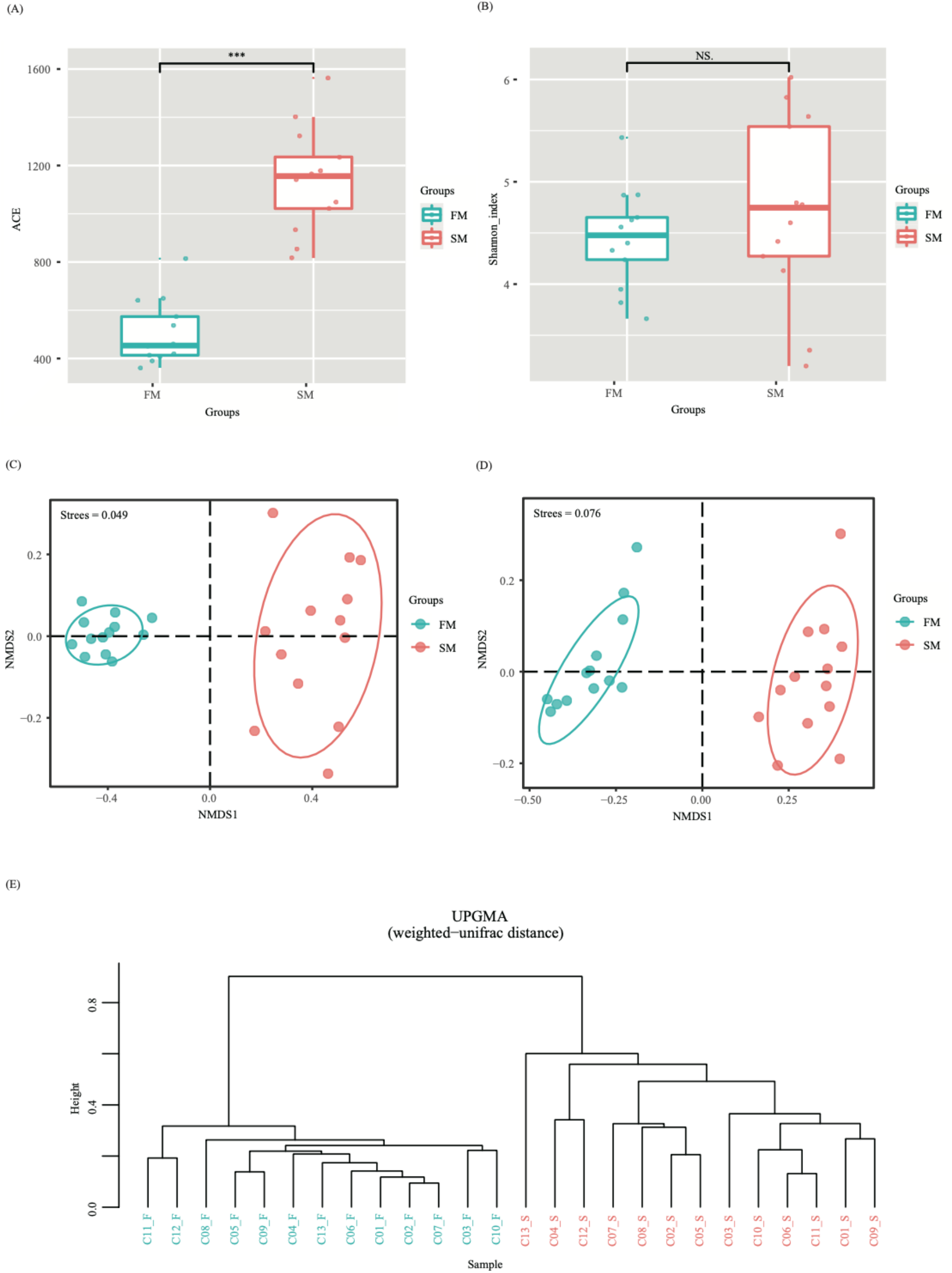
Diversity analysis of gut (FM) and skin (SM) microbial composition in larvae of *I. bannanicus* (*N* = 13). (A) ACE index; (B) Shannon index. The bottom and top of the box are the first and third quartiles, the band inside the box is the median, and the ends of the whiskers represent the minimum and maximum. Asterisk indicates significant difference (*P* < 0.001, paired Wilcox test). NS indicates no significant difference between groups. (C) Non-metric multidimensional scaling (NMDS) analysis with weighed unifrac distance (R = 0.99; *P* = 0.001; ANOISM test was performed) showing bacterial composition across gut (FM) and skin (SM) samples. (D) Non-metric multidimensional scaling (NMDS) analysis with unweighed unifrac distance (R = 1; *P* = 0.001; ANOISM test was performed) showing bacterial composition across gut (FM) and skin (SM) samples. Each point in the graph represents a sample, the distance between the points indicate the degree of difference. Samples of the same group are represented in the same colour. Each group adds to the 80% confidence ellipse in NMDS analysis. (E) The UPGMA cluster analysis of gut (C01_F-C13_F and skin (C01_S-C13_S) samples in weighed Unifrac distances. The figure represents UPGMA cluster tree.

To reveal compositional change we conducted Non-metric multidimensional scaling analysis (NMDS) between the gut and skin samples (Fig. 2C; Fig 2D) based on the OTUs and found both samples clustered separately for both weighed unifrac distance (R = 0.99; *P* = 0.001; Fig. S4C) as well as for the unweighed unifrac distance (R = 1; *P* = 0.001; Fig. S4D).

### Microbial diversity across gut and skin samples of *I. bannanicus*

Results for the UPGMA weighed unifrac distance analysis showed that relative abundance and distribution of bacteria may be indicative of variation in the samples of gut and skin microbiome composition (Fig. 2E).

### Unique and shared microbial taxa

The indicator OTU concept provided a framework for investigating the differences of microbial communities in the two groups (gut and skin) at species level. The gut samples shared exclusively 32 OTUs in common (Fig. 3A) at species level. Further, 61 OTUs were exclusively shared in the skin samples shared (Fig. 3B) at species level. Results obtained by conducting t-test (Table S3; Fig. 3C) revealed significant differences in the microbial diversity between gut and skin samples at phylum level. The skin samples had higher diversity when compared to the gut samples (Fig. S4A). The gut and skin samples did not share any unique OTUs in common at species level (Fig. S4B).

**Figure 3.**
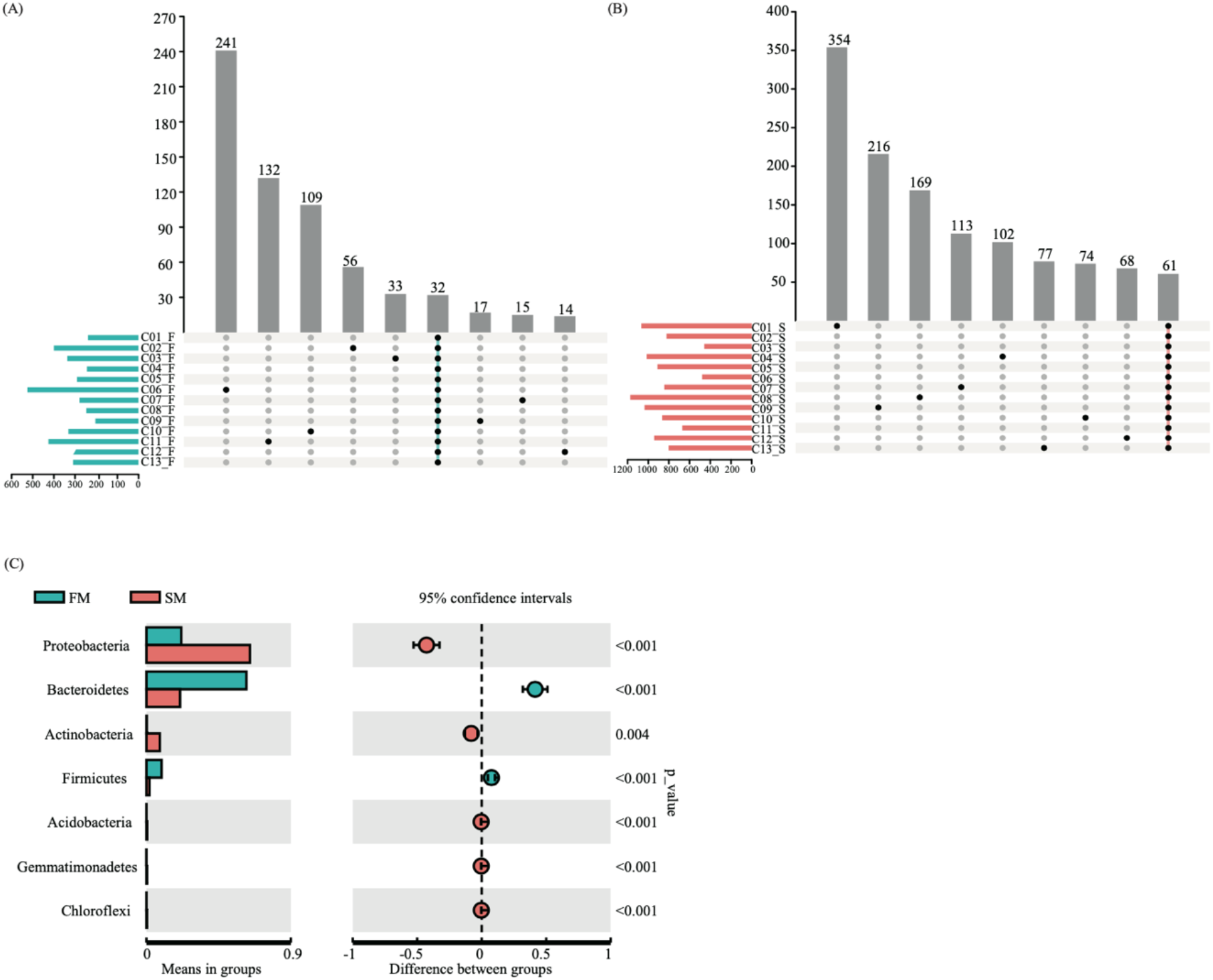
(A) Shared bacteria in the gut samples (C01_F-C13_F) of *I. bannanicus* larvae; (B) Shared bacteria in the skin samples (C01_S-C13_S) of *I. bannanicus* larvae; (C) Results of t-test between gut (FM) and skin (SM) samples of *I. bannanicus*. Each bar represents mean value of the phyla diversity with significant differences (*P* < 0.005) in the abundance between groups.

### Taxonomic and functional characteristics of bacteria between the gut and skin microbiota

We investigated differences in the gut and skin samples of *I. bannanicus* larvae at OTU level. We analysed the enrichment according to their taxonomy using Manhattan plot (Fig. 4). In both samples, OTUs enriched in *I. bannanicus* belonged to wide range of bacterial phyla including Acidobacteria, Actinobacteria, Bacteriodetes, Chloroflexi, Cyanobacteria, Firmicutes, Planctomycetes, Proteobacteria and Verrucomicrobia (False Discovery Rate (FDR) adjusted *P* < 0.05, Wilcoxon rank-sum test; Fig. 4).

**Figure 4.**
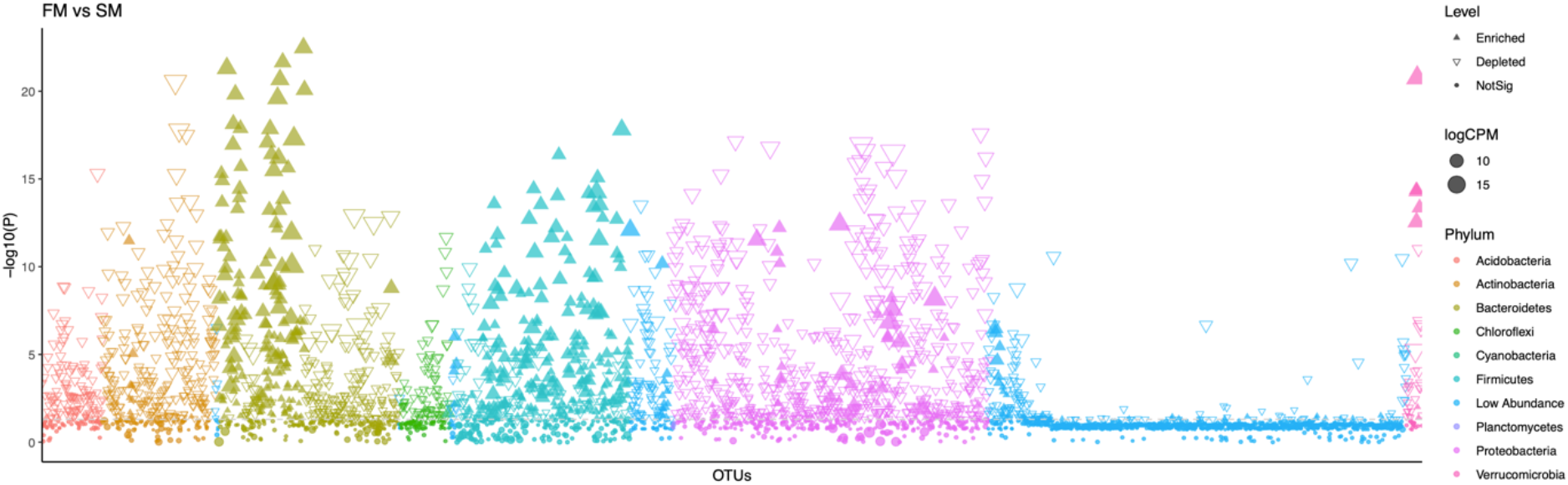
Taxonomic and functional characteristics of differential bacteria between the gut and skin microbiota in larvae of *Ichthyophis bannanicus*. Manhattan plot showing OTUs enriched in gut and skin. Each dot or triangle represents a single OTU. OTUs enriched in gut and skin are represented by filled or empty triangles, respectively (False discovery rate (FDR) adjusted *P <* 0.05, Wilcoxon rank-sum test). OTUs are arranged in taxonomic order and coloured according to the phylum. Counts per million reads mapped (CMP).

### Linear discriminant analysis effect size (LEfSe)

We compared the compositional similarity between the gut and skin samples of *I. bannanicus* larvae by calculating the pairwise distance among OTU abundance (Fig. 5A). The LEfSe analysis was used to analyse the species abundance between the gut and skin samples in *I. bannanicus*, which showed that there are 52 biomarkers with an LDA score > 4. A cladogram representing microbial taxa enriched in gut (red colour) versus skin (green colour) samples of *I. bannanicus* is presented in Fig. 5B.

**Figure 5.**
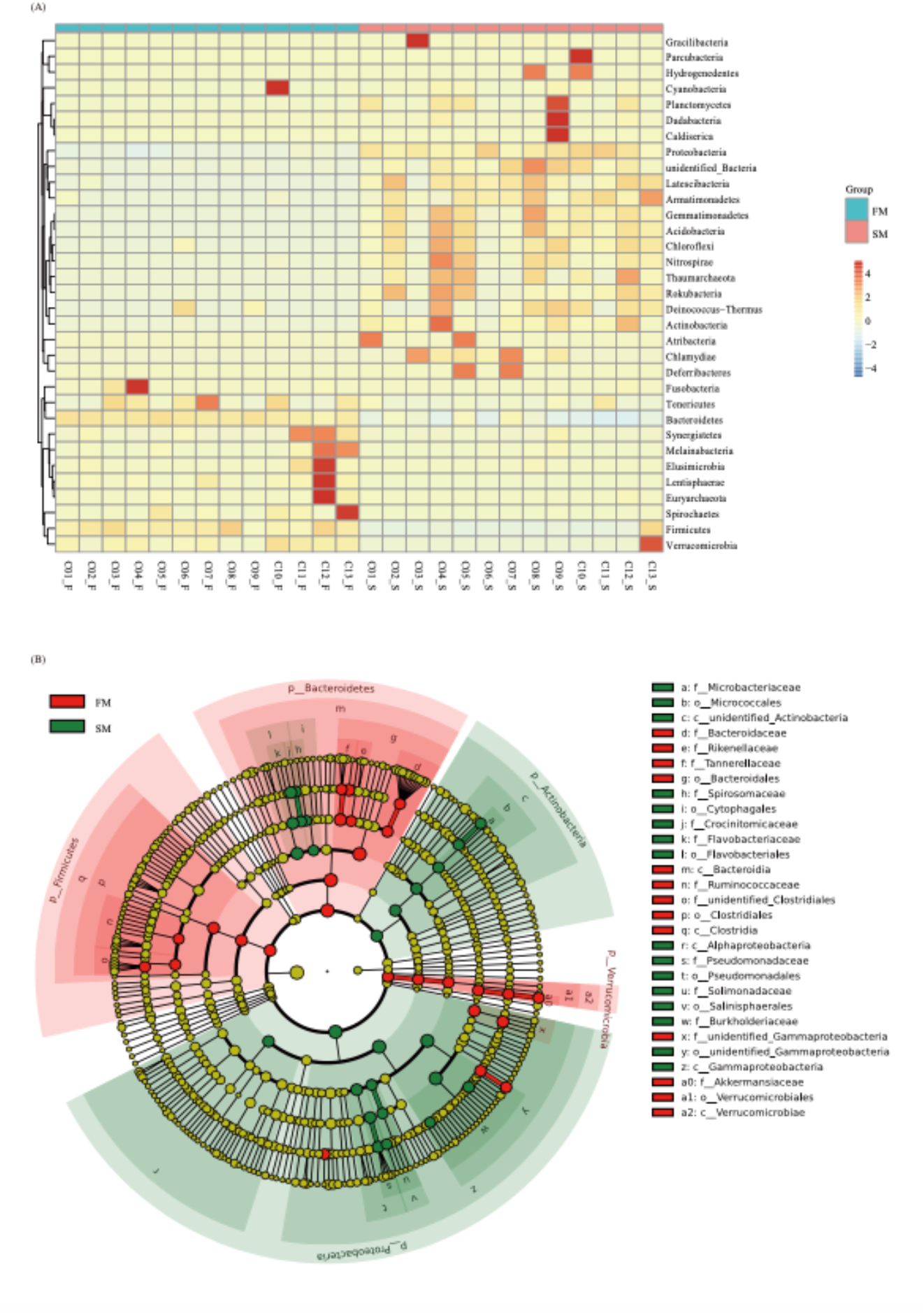
(A) A clustered heatmap illustrating top 33 mean abundances of the bacterial community taxa assigned to phyla level. The colour scale of higher (red) and lower (blue) shows the relative abundances of bacterial communities in the gut (C01_F-C13_F) and skin (C01_S-C13_S) samples; (B) Linear discriminatory analyses (LEfSe) of bacterial taxa. LDA value distribution clade map shows abundance of OTUs in larvae of *I. bannanicus* according to gut (FM) and skin (SM) samples (Biomarkers) with LDA score > 4. Clade map shows classification level from phylum to genus (circles radiating from inside to outside). Each small circle at a different classification level represents a classification at that level, and the diameter of the small circle is proportional to the relative abundance in larvae of *I. bannanicus* according to gut and skin samples. The species with no significant difference is uniformly colored yellow and the different species Biomarker follow group colors. The red node indicates the microbial group that plays an important role in the gut samples and the green node indicates the important role in the skin samples. If the microbial groups are missing, it indicates that there is no significant difference in that particular group.

### Microbial interaction in the gut samples of *I. bannanicus*

The 16S rRNA sequencing data was used to create microbial interactions in the gut samples, which provided potential interaction patterns of microbes (Fig. 6). The network map created for bacterial interactions showed nodes and connections as presented in Fig. 6 (network diameter = 15; modularity = 0.521; clustering coefficient = 0.484; graph density = 0.0316; average degree = 11.827; average path length = 5.085).

**Figure 6.**
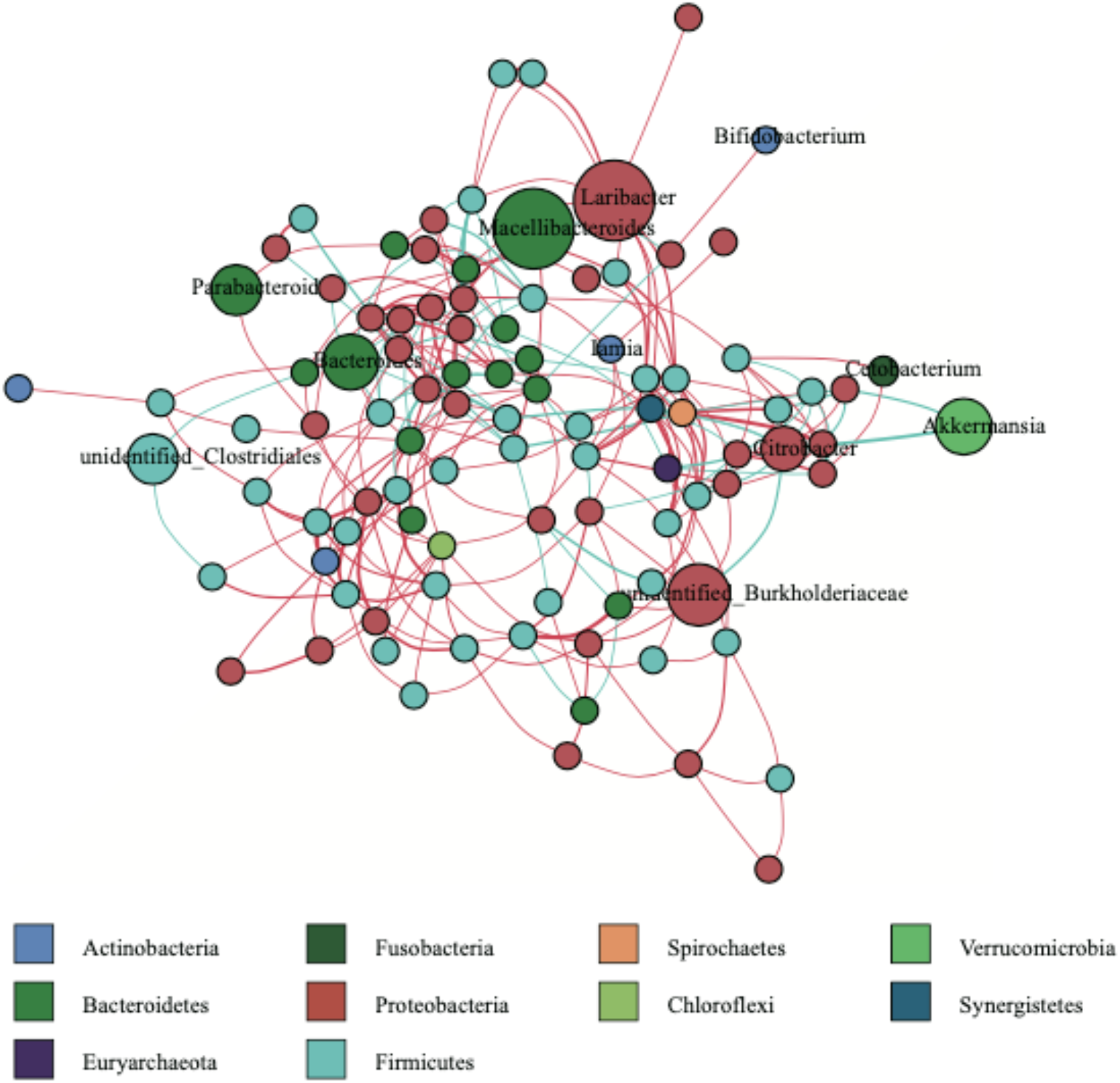
Significant co-occurrence and co-exclusion relationships among the abundances of clades in the gut samples (*N* = 13) *I. bannanicus* larvae. Each node represents average relative abundance of the bacterial genera. Nodes of the same phylum have same colour. The thickness of line between the nodes and the correlation coefficient of species interaction are positive in the absolute value. Red colour connections correspond to the positive correlation while blue colour connections correspond to negative correlation.

## DISCUSSION

The advent of next-generation sequencing technologies (NGS) has helped unravel the complex host-microbial interactions, especially in terms of estimating microbial diversity. The older culture-dependent methods, which are tedious to implement, are being replaced by culture-independent ones, leading to a rapid accumulation of data on microbiome diversity. Our analysis of the BM of caecilians too, has been made possible through this.

Amphibian skin provides a suitable environment for rich communities of microorganisms to flourish. These play an important role as symbionts that aid in protection (pathogens, diseases, infections) and physiological functions such as electrolyte exchange and respiration. The bacteria found in the gut contribute to the nutritive functions. The central idea of our study was to characterize the BM of *I. bannanicus* larvae, which had previously not been studied. As the larvae of caecilians are unique among amphibians in being carnivorous, we hypothesized that the gut microbiota of *I. bannanicus* would be distinct at a higher taxonomic level. Our study shows that some of the unspecialized bacterial species are shared between the gut and skin. This is probably aided by the aquatic medium that they inhabit. A study of redback salamanders showed that the bacterium *Janthinobacterium lividum*, which plays a protective role, was able to survive passage through the gut, suggesting that the gut could act as a reservoir for protective skin bacteria (*Wiggins et al., 2011*). The skin BM is directly influenced by the external environment, which the gut BM is mediated by internal factors that aid in digestion. Thus, the gut and skin have different influencing gradients that determine the colonization and re-colonization of the BMs.

Members of Actinobacteria, Bacteriodetes, Firmicutes and Verrucomicrobia are seen to be shared in all gut samples of *I. bannanicus* larvae. In total 32 OTUs occur in common in all gut samples. Members of phylum Actinobacteria, Bacteriodetes, Chloroflexi, Gemmatimonadetes and Proteobacteria are seen to be shared in all skin samples of *I. bannanicus* larvae. In total 61 OTUs seen to co-occur in all skin samples. Our analysis shows that, bacterial diversity (at species level) was high on skin when compared to the gut samples. Also, skin and gut seem to harbor unique specialized bacteria that may be essential to perform certain functions. Few species of bacteria were seen to co-occur in both gut and skin samples; these species might be part of the gut-skin axis in larvae of *I. bannanicus*, which we consider to be unspecialized in function. Results of our study suggests that skin, acting in the capacity of not only a barrier but also as a respiratory surface, is more influenced by external abiotic factors when compared to gut, where bacterial diversity is higher when compared to the gut BM.

By consuming oxygen and lowering redox potential in the gut environment, proteobacteria are thought to play a vital role in preparing the gut for colonization of anaerobes that are required for healthy gut function (*Moon et al., 2018*). Proteobacteria and Actinobacteria strongly affect normal microbiota composition. Actinobacteria play an important role in the decomposition of organic materials. Bacteriodetes are known contribute to the preservation of gut microbalance, immune system development, polysaccharide degradation, and nutritional use acceleration (*Tong et al., 2019a*). Firmicutes aid in digestion, support immune functions and also influence behaviour in animals. Firmicutes help in carbohydrate fermentation and nutrient absorption (*Tong et al., 2019a*). Previous studies report that the Firmicutes - Bacteriodetes ratio indicates a higher efficiency of energy uptake from food (*Tong et al., 2019a*), which thereby enhances fitness. Microbes belonging to the Verrucomicrobia are mucin-degrading bacteria that play a major role in glucose homeostasis and contribute to intestinal health.

Comparative fecal sample analysis conducted between larvae belonging to amphibian orders showed that in case of anuran tadpoles, the most dominant gut bacterial phyla were Proteobacteria > Fusobacteria > Firmicutes > Bacteriodetes > Cyanobacteria, whereas in case of caecilian larvae the relative dominance of bacterial phyla was Bacteriodetes > Proteobacteria > Firmicutes > Verrucomicrobia > Actinobacteria. Further, in the case of salamander larvae, the most dominant bacterial phyla were Proteobacteria > Bacteriodetes > Firmicutes > Actinobacteria > Verrucomicrobia. Anurans constitute predominantly of Cyanobacteria, whereas in case of caecilians and salamanders (*Sanchez et al., 2017*), Verrucomicrobia and Actinobacteria were represented (comparison made only among the most dominant five phyla). Our study shows that though the bacterial diversity overlaps to some extent, the species abundance of dominant phyla was different. The similarities between the caecilian and salamander microbiota can be corelated with the aquatic phase of their life history stage. Unlike in other amphibians, caecilian skin is highly glandular and secretes substances that are important for chemical defense against predators and microorganisms (*Duellman & Trueb, 1986; Jared et al., 1999*).

The data presented here are based on *ex situ* studies. It is not yet known whether the skin and gut microbiomes of larval *I. bannanicus* inhabiting natural environments would also comprise of above-mentioned bacterial phyla. We assume that the caecilian larvae would also have similar core microbiomes, as shown to be the case in a study conducted in fire salamanders (*Demircan et al., 2018*). Diet preferences in wild, for the caecilian larvae may have differential prey choices including the Chironomous larvae. Thus, to some extent the gut microbiome would have more diversity than the laboratory reared individuals. The caecilian larvae inhabiting in natural water bodies are exposed to various hetero-specific individuals, contributing to the additional microbes on skin. Although the bacterial microbiomes occurring in gut and skin of caecilian larvae in wild may vary to some extent than our laboratory based study, organisms are now known to maintain a core microbiome despite changes in environment (*Tong et al., 2019a*). In our study, all the test subjects survived the period of experimentation, during which no morbidity was observed.

## CONCLUSION

Microbes colonize virtually all epithelial surfaces, lumen and mucosa of organisms, wherein they often outnumber the somatic cells of their hosts, implying the importance of microorganisms in host physiology (*Hanning & Diaz-Sanchez, 2015; Davenport et al., 2017*). The present study contributes to the bacterial characterization of gut and skin of *I. bannanicus* using 16S rRNA gene amplicon sequencing. The study provides a comprehensive account of the gut and skin BM of *I. bannanicus*, the only caecilian species in China. Our study reveals that though the bacterial diversity of the gut partially overlaps among larvae of the three amphibian orders, the relative abundance of the dominant phyla remains distinct. We also show that the skin BM is more diverse than the gut BM. The findings suggest that specific microbes play an important role in an individual, which promotes metabolic flexibility, adaptation to the changing environment, protection from diseases and infections. Thus, understanding the microbial communities and describing the local bacterial communities associated with gut and skin of *I. bannanicus* larvae is important. The knowledge generated by this study forms a foundation for further exploration of the BM of Gymnophiona and facilitates comparisons of the BMs of larvae across the amphibian orders.

## Acknowledgements

We thank Cheng-Hai Fu for the maintenance of animals during the study.

## Funding

This study was financially supported by the funding from Guangxi University Special Talent Recruitment Grant to Madhava Meegaskumbura. This study was also supported by the Postdoctoral Project from Guangxi University to Amrapali Prithvisingh Rajput.

## Competing interests

The authors declare that the research was conducted in the absence of any commercial or financial relationships that could be construed as a potential conflict of interest.

## Author contributions

- Amrapali Prithvisingh Rajput conceived and designed the experiments, performed the experiments, analyzed the data, wrote the manuscript with revisions from Madhava Meegaskumbura, prepared figures and tables, and drafted the successive versions of this paper.
- Zhou Shipeng made figures, reviewed and edited the draft.
- Madhava Meegaskumbura conceived and designed the experiments, authored or reviewed drafts of the paper and approved the final draft.

## Animal ethics

This study was carried out in accordance with the recommendations of Institutional Animal Care and Use Committee of Guangxi University (GXU), Nanning-China. Animal procedures were approved by GXU (GXU2019-071).

## Data availability

All data pertaining to the study will be made available on GenBank (upon acceptance).

## Notes

### Competing Interest Statement

The authors have declared no competing interest.

